# Temporal identity establishes columnar neuron morphology, connectivity, and function in a *Drosophila* navigation circuit

**DOI:** 10.1101/485185

**Authors:** Luis F. Sullivan, Timothy L. Warren, Chris Q. Doe

**Affiliations:** Institute of Neuroscience, Institute of Molecular Biology, Howard Hughes Medical Institute, University of Oregon, Eugene, OR 97403

**Keywords:** *Drosophila*, neuroblasts, intermediate neural progenitors, temporal patterning, synaptic connectivity, central complex, ellipsoid body, protocerebral bridge, Eyeless, celestial navigation

## Abstract

The insect central complex (CX) is a conserved brain region containing 60+ neuronal subtypes, several of which contribute to navigation. It is not known how CX neuronal diversity is generated or how developmental origin of subtypes relates to function. We mapped the developmental origin of four key CX subtypes and found that neurons with similar origin have matching axon/dendrite targeting. Moreover, we found that the temporal transcription factor (TTF) Eyeless/Pax6 regulates the development of two recurrently-connected CX subtypes: Eyeless loss simultaneously produces ectopic P-EN neurons with normal axon/dendrite projections, and reduces the number of E-PG neurons. Furthermore, the transient loss of Eyeless during development impairs adult flies’ capacity to perform celestial navigation. We conclude that neurons with similar developmental origin have similar connectivity, that Eyeless maintains equal E-PG and P-EN neuron number, and that Eyeless is required for the development of circuits that control adult navigation.

## Introduction

Work over the past two decades has revealed two important developmental mechanisms that generate neuronal diversity from flies to mice. First, spatial patterning cues produce different pools of neural progenitors (called neuroblasts in insects); second, neuronal progenitors/neuroblasts sequentially express a series of transcription factors that generate additional neuronal diversity (Kohwi and Doe 2013). These so-called “temporal transcription factors” or TTFs are expressed transiently in progenitors, are inherited by neurons born during the expression window, and specify progenitor-specific neuronal identity (Rossi et al. 2016; Doe 2017). For example, the Hunchback (Hb) TTF is present in *Drosophila* embryonic neuroblasts as they produce their first progeny; loss of Hb leads to absence of first-born neurons, whereas prolonging Hb expression generates ectopic first-born neurons (Isshiki et al. 2001). While TTFs are clearly important for generating molecularly distinct neuronal subtypes, their role in establishing neuronal morphology, connectivity, and behavior remains relatively poorly understood.

Recent work has identified candidate TTFs expressed during larval brain neuroblast lineages. In particular, there are four bilateral “type II” neuroblasts, DM1-DM4, that generate most of the intrinsic neurons of the central complex (CX) (Figure 1A) (Yang et al. 2013). Type II neuroblasts have a complex lineage. They produce a series of molecularly distinct intermediate neural progenitors (INPs), which in turn each produce 4-6 molecularly distinct ganglion mother cells (GMCs) that yield a pair of sibling neurons (Figure 1B) (Bello et al. 2008; Boone and Doe 2008; Bowman et al. 2008). Several laboratories have identified candidate temporal transcription factors (TTFs) that are sequentially expressed in type II neuroblasts (Figure 1B, horizontal axis) or INPs (Figure 1B, vertical axis) (Bayraktar and Doe 2013; Ren et al. 2017; Syed et al. 2017).

**Figure 1.**
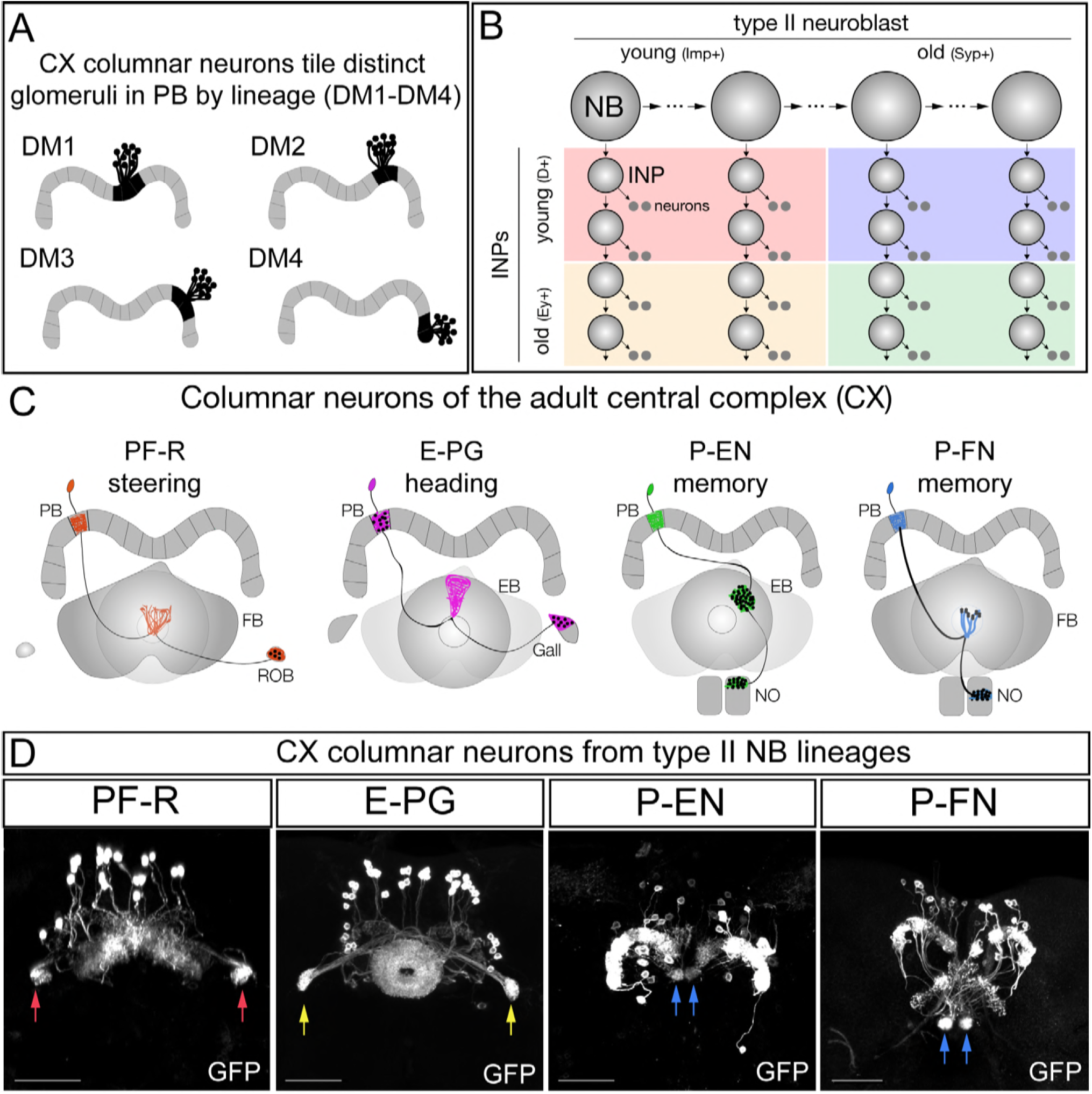
CX columnar neurons are generated by type II neuroblast lineages. (A) CX columnar neurons originate from four bilateral type II neuroblast lineages (DM1-DM4), with DM1 lineage neurons innervating the most medial PB glomeruli, and DM4 lineage neurons innervating the most lateral PB glomeruli. Adult brain right hemisphere shown. (B) Type II neuroblasts and their INP progeny both express temporal transcription factors that subdivide the lineage into distinct developmental windows. Finer subdivisions exist but are not shown for clarity. (C)PF-R, E-PG, P-EN, and P-FN columnar neuron subtypes; each has a proposed function in navigation (line 2) and a distinct pattern of connectivity. (D) Adult CX columnar neurons derived from INPs labeled with adult LexA lines specific for each subtype; see Figure 1 – supplement 1A for genetic details. ROB, red arrows; Gall, yellow arrows; Noduli, blue arrow. Scale bars, 40μm

In this study we address how larval brain TTFs contribute to the development and function of the adult insect central complex (CX). The CX is a highly conserved brain region in insects that is thought to play a crucial role in navigation and motor control (Pfeiffer and Homberg 2014; Green et al. 2017; Heinze 2017; Kim et al. 2017; Franconville et al. 2018; Green et al. 2018). The CX is characterized by four distinct neuropil regions: the Ellipsoid Body (EB), Fan-shaped Body (FB), Protocerebral Bridge (PB), and Noduli (NO); the CX is also connected to lateral neuropils termed the Gall and the Round body (ROB) (Wolff et al. 2015). Columnar neurons, which innervate single glomeruli that tile the entire EB and PB neuropil, have been shown to play a key role in navigation (Pfeiffer and Homberg 2014; Green et al. 2017; Heinze 2017; Kim et al. 2017; Turner-Evans et al. 2017; Franconville et al. 2018; Green et al. 2018). There are at least four columnar neuron subtypes (Figure 1C). The E-PG neurons have spiny dendritic arbors in the EB (hence the E at the front of their name) and provide outputs to the PB and Gall (hence the PG at the end of their name); conversely, P-EN neurons have spiny dendritic arbors in the PB and provide outputs to the EB and Noduli. Recently it has been proposed that the E-PG/P-EN neurons form a recurrent circuit that tracks the fly’s orientation in space (Lin et al. 2013; Green et al. 2017; Turner-Evans et al. 2017; Green et al. 2018). Two additional columnar neuron classes are PF-R neurons that have dendritic spines in the PB and FB and project axons to the ROB, and the P-FN neurons which have dendritic spines in the PB and project axons to the FB and Noduli (Figure 1A) (Wolff et al. 2015; Wolff and Rubin 2018); both are proposed to have a role in navigation based on anatomical connectivity (Heinze 2017; Wolff and Rubin 2018), but their function has not been experimentally determined.

Here we map the developmental origin of these four CX neuronal subtypes postulated to have a critical role in navigation. We find that each is derived from a specific temporal window during the INP cell lineage, and that neurons with similar developmental origins have similar axon/dendrite neuropil targets. We show that the TTF Eyeless, expressed in the latter half of INP lineages, is required both to promote the identity of the two CX neuron subtypes born late in INP lineages (E-PG, PF-R) as well as to repress the identity of two CX neuron subtypes born during early INP lineages (P-EN, P-FN). In this way, the Eyeless TTF regulates the relative proportion of each neuronal subtype: loss of Eyeless generates fewer E-PG neurons and more P-EN neurons. Importantly, the ectopic P-EN neurons have normal anatomical connectivity. Finally, we show that loss of Eyeless specifically during the larval stages when E-PG neurons are born results in a highly specific defect in adult flight navigation, consistent with the proposed role of E-PGs in maintaining an arbitrary heading to a sun stimulus. Our findings are the first to identify the developmental origin of functionally important adult flight navigation neurons. Moreover, they set the stage for manipulating developmental genetic programs to alter the number and function of each class of adult CX neurons.

## Results

### CX columnar neurons are generated by type II neuroblast lineages

We used intersectional genetics to map the developmental origin of four CX columnar types (Figure 1 – Supplement 1). Our strategy was to use the FLP enzyme to permanently open a *lexAop-FRT-stop-FRT-GFP* reporter in specific populations of INPs and then use adult columnar neuron LexA transgenes to determine the number of each adult columnar neuron type made by each of these INP populations. This approach allowed us to map the developmental origin of neurons labeled by LexA reporters only at pupal or adult stages. We opened the *lexAop-FRT-stop-FRT-GFP* reporter in all INPs of the type II neuroblast lineages and confirmed that all four types of adult CX columnar neurons are generated by type II neuroblasts (Figure 1 – Supplement 1A). Indeed, we found that type II neuroblasts make all 30 PF-R neurons, all 40 E-PG neurons, all 40 P-EN neurons, and all 50 P-FN neurons (Figure 1D). We conclude that the four types of CX columnar neurons are all derived from type II neuroblast lineages.

### CX columnar neurons are generated by young type II neuroblast lineages

Larval type II neuroblasts produce neurons over five days (0-120h after larval hatching; ALH). We used intersectional genetics to determine when each columnar neuron subtype was born during the type II neuroblast lineage. We transiently expressed the FLP recombinase in INPs to permanently open the lexAop reporter at different times during type II neuroblast lineages and assayed for the number of PF-R, E-PG, P-EN, or P-FN adult neurons made at each time-point (method summarized in Figure 1 – Supplement 1B). We found that PF-R neurons were made first in larval type II neuroblast lineages, followed by E-PG neurons, and then by P-EN and P-FN neurons which share overlapping birthdates (Figure 2A). The relatively broad distribution of columnar neuron birthdates is likely due to DM1-DM4 individual lineages generating neuron subtypes asynchronously, but could also represent natural developmental variation or stochasticity in the time of columnar neuron birthdates, but it is most consistent with each pool of 30-50 columnar neurons being generated within a 12h temporal window in the type II neuroblast lineage (Figure 2B). We conclude that all four types of columnar neurons arise early in type II lineages.

**Figure 2.**
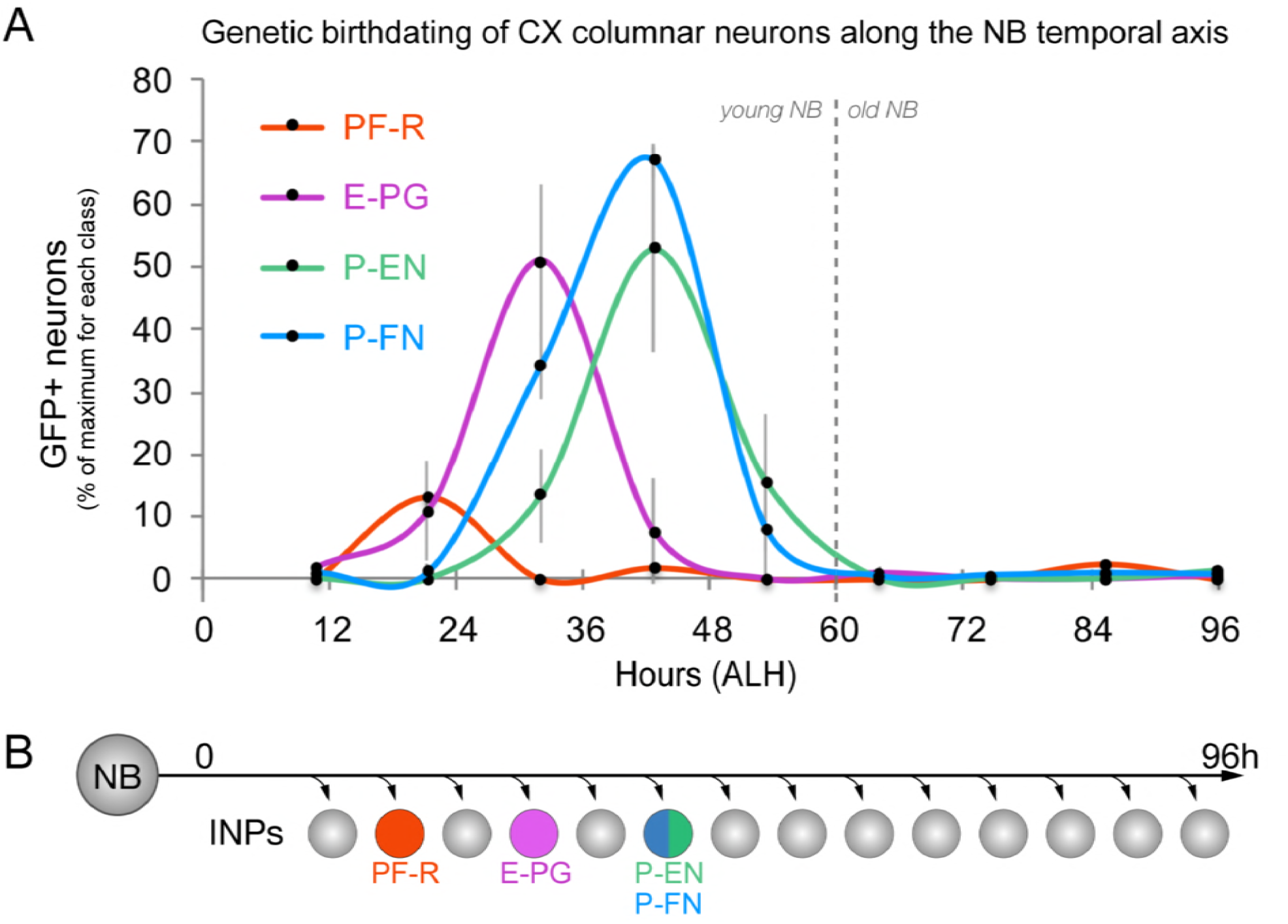
CX columnar neurons are generated by young type II neuroblast lineages. (A) Identifying the time during the neuroblast lineage that produces each columnar neuron subtype. See Figure 1 – supplement 1B for genetic details. Note that PF-R neurons are born first, E-PG neurons second, and then P-EN/P-FN neurons sharing a common birthdate (n=3-6 per time-point). (B) Summary of NB birthdating results.

### CX columnar neurons with similar developmental origin have similar axon/dendrite targeting

We next defined columnar neuron birthdates along the INP temporal axis (see Figure 1 – Supplement 1C). Young INPs express Sox family transcription factor Dichaete (D), whereas old INPs express the Pax6 family transcription factor Eyeless (Bayraktar and Doe 2013; Eroglu et al. 2014; Farnsworth et al. 2015). Here we test whether columnar neuron subtypes arise from a young D+ or old Ey+ temporal window. As expected, all columnar neuron subtypes are labeled when the lexAop reporter is ‘opened’ in all INPs (Figure 3A-D). In contrast, when the lexAop reporter is ‘opened’ only in old INPs, we detect all 40 E-PG and all 30 PF-R adult neurons but no P-EN or P- FN neurons (Figure 3E-H). We conclude that all P-EN and P-FN neurons are born from young INP lineages, whereas all E-PG and PF-R neurons are born from old INP lineages (summarized in Figure 3I). Interestingly, the P-EN and P-FN columnar neurons have a highly similar developmental origin and project to similar CX neuropils (dendrites to PB, axons to Noduli; Figure 3I), whereas E- PG and PF-R columnar neurons have distinct developmental origins and share no similarities in neuropil targets, suggesting that developmental origin may be tightly linked to neuronal morphology and anatomical connectivity (see Discussion).

**Figure 3.**
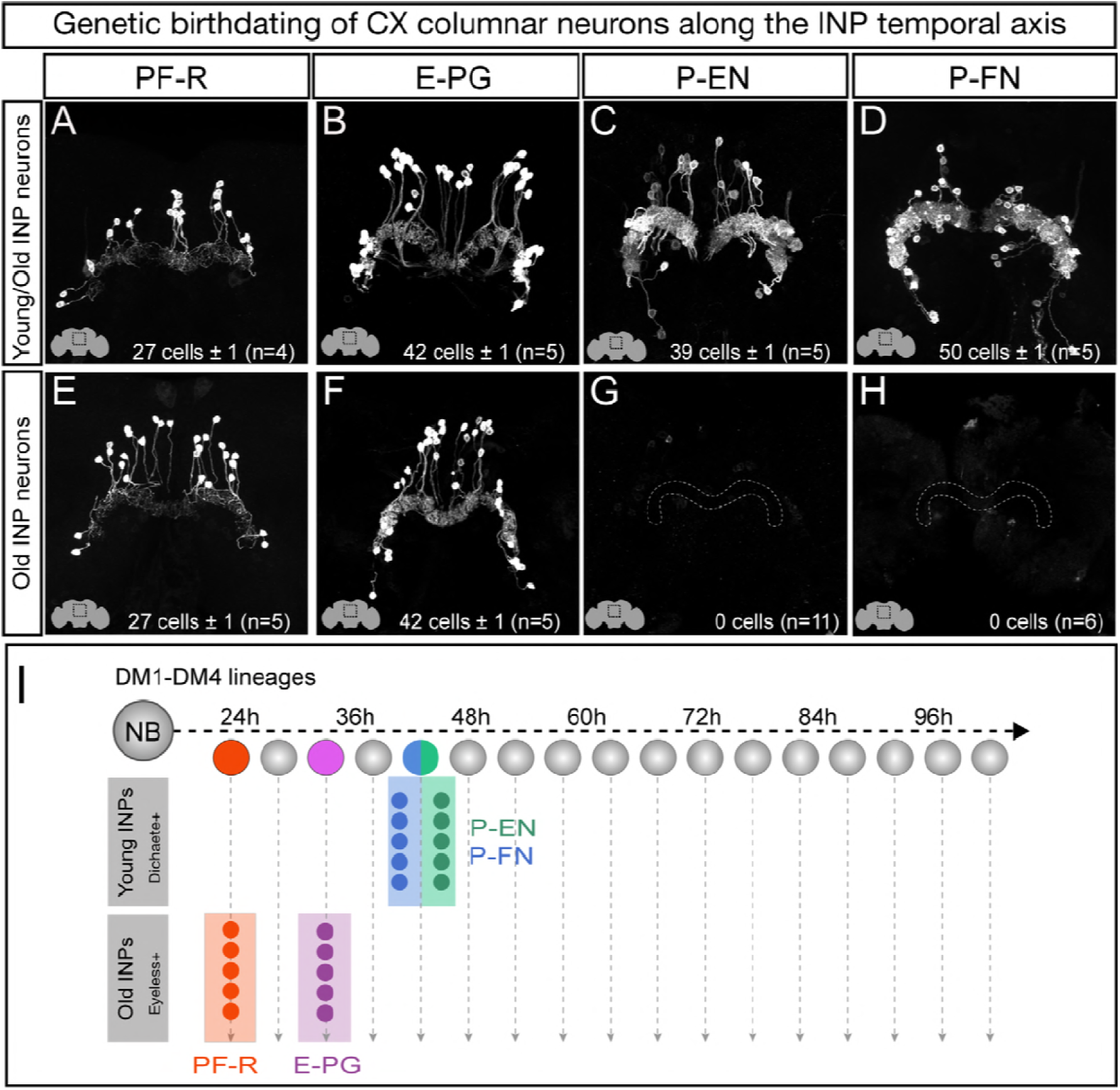
Each CX columnar neuron type arises exclusively from young or old INP lineages. (A-D) All columnar neurons labeled by subtype-specific LexA lines derive from INP lineages (n=5 for each experiment). See Figure 1 – supplement 1A for genetic details. (E-H) The PF-R and E-PG columnar neurons are generated by late INPs (n=5 for each experiment), whereas the P-EN (n=11) or P-FN (n=6) neurons were not derived from old INPs and thus are fully derived from young INPs. See Figure 1 – supplement 1C for genetic details (I) Summary of INP birthdating results.

### The Eyeless temporal transcription factor promotes E-PG and PF-R molecular identity

Our birthdating results indicated that INP age might be a major determinant of CX columnar neuron morphology and connectivity. We next tested whether the TTF Eyeless, which is expressed by INPs during the last half their lineage, specifies the identity of PF-R and E-PG neurons, which are born from Ey+ INPs. To knock down Eyeless expression in INPs, we used an *eyeless* enhancer-Gal4 line (*R16B06-Gal4*) to drive a *UAS-Ey*^*RNAi*^ transgene that we previously showed eliminates all detectable Eyeless protein (Bayraktar and Doe 2013).

In wild type adults, there are ∼40 E-PG neurons and ∼30 PF-R neurons (Figure 4A,B; quantified in G,H). In adults where Ey^RNAi^ is expressed in old INPs, we found nearly complete loss of PF-R and E-PG neurons (Figure 4D,E; quantified in G,H); we suggest that these neurons are transformed into an early-born INP progeny identity, but we can’t rule out that they undergo apoptosis. In addition, we performed an antibody screen for neuronal markers of CX neuronal subtypes, and identified Toy as specifically marking all PF-R and E-PG neurons but none of the young INP-derived P-EN and P-FN neurons (Bayraktar and Doe 2013). Here we show that Toy+ neurons generated by old INPs are also significantly reduced following Ey^RNAi^ in old INPs (Figure 4C,F; quantified in I). We conclude that the Eyeless temporal transcription factor is required for the specification of PF-R and E-PG columnar neurons.

**Figure 4.**
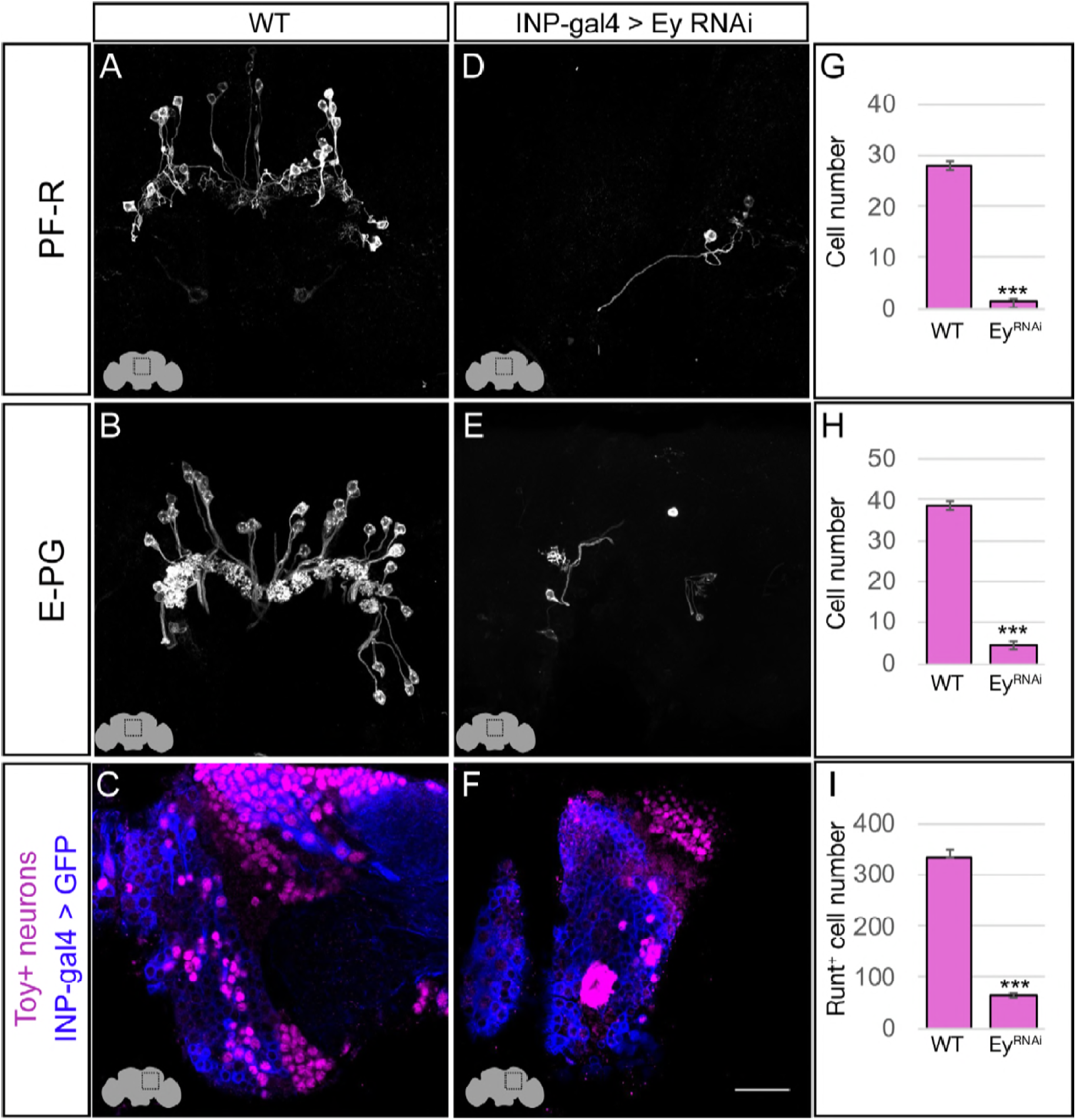
Eyeless promotes PF-R and E-PG molecular identity. (A-C) Wild-type numbers of PF-R, E-PG, and Toy^+^ neurons in the dorsoposterior adult brain. See methods for genotypes. (D-F) Eyeless^RNAi^ in INP lineages decreases the number of PF-R, E-PG, and Toy^+^ late-born neurons in the dorsoposterior adult brain. See methods for genotypes. (G-I) Quantification (n=5 for each experiment). ***, p <0.001. Scale bar, 20μm

### The Eyeless temporal transcription factor represses P-EN and P-FN molecular identity

The P-EN and P-FN columnar neurons derive from early INP progeny, prior to the expression of Eyeless in later-born INPs, raising the question of whether Eyeless expression triggers a switch from early-born P-EN/P-FN production to late-born E-PG/PF-R production. To determine if Eyeless terminates production of early-born P-EN and P-FN columnar neurons, we expressed Ey^RNAi^ in old INPs, and assayed for ectopic P-EN or P-FN neurons. In wild type adults, there are ∼40 P-EN neurons and ∼50 P-FN neurons (Figure 5A,B; quantified in G,H). In adults where Ey^RNAi^ was expressed in old INPs, we found an over two-fold increase in the number of P-EN and P-FN neurons (Figure 5D,E; quantified in G,H). In addition, the antibody screen described above identified the transcription factor Runt as specifically marking all early-born P-EN and P-FN neurons but none of the late-born E-PG and PF-R neurons (data not shown). In wild type, there are ∼220 Runt+ adult neurons made by INP progeny, but Ey^RNAi^ led to a significant increase to ∼580 Runt+ adult neurons (Figure 5C,F; quantified in I), consistent with a role for Eyeless in terminating production of young INP-derived neurons. We conclude that Eyeless maintains equal pools of E-PG and P-EN neurons by triggering a switch from early-born P-EN/P-FN neurons to late-born E-PG/PF- R neurons.

**Figure 5.**
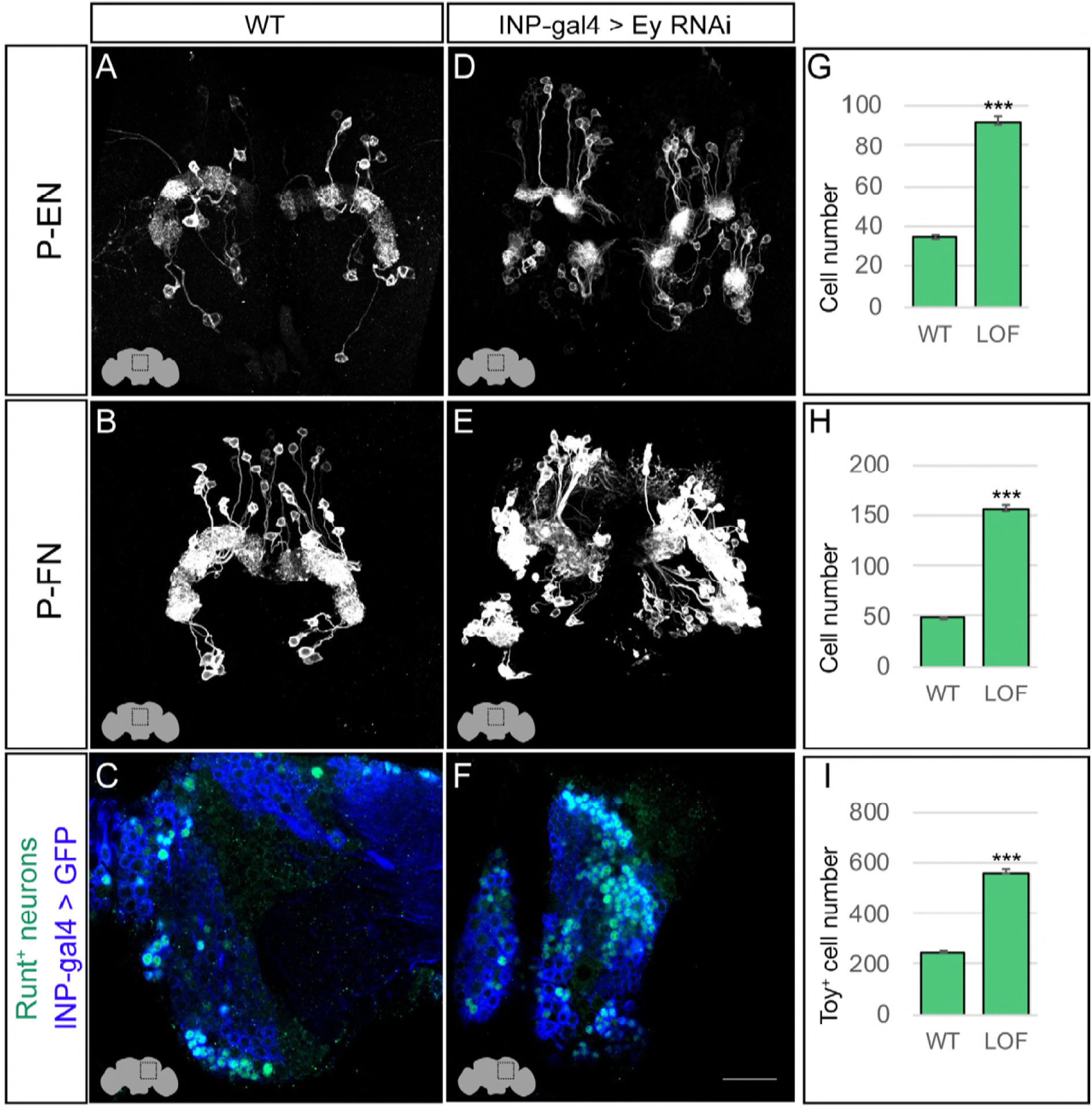
Eyeless represses P-EN and P-FN molecular identity. (A-C) Wild-type numbers of P-EN, P-FN, and Runt^+^neurons in the dorsoposterior adult brain. See methods for genotypes. (D-F) Eyeless^RNAi^ in INP lineages increases the number of P-EN, P-FN, and Runt^+^ late-born neurons in the dorsoposterior adult brain. See methods for genotypes. (G-I) Quantification (n=5 for each experiment). ***, p <0.001. Scale bar, 20μm

### Loss of Eyeless produces ectopic P-EN neurons with normal morphology and anatomical connectivity

Loss of Eyeless extends the production of P-EN neurons into an older stage of INP lineages, creating a mismatch between their molecular temporal identity (early) and their time of differentiation (late). We tested whether the ectopic P-EN neurons have a neuronal morphology and anatomical connectivity characteristic of the endogenous early-born neurons, or whether their later birthdate results in different morphology or connectivity. We designed a genetic method for specifically labeling the ectopic late-born P-EN neurons – but not the endogenous early-born P-EN neurons – to trace their morphology and anatomical connectivity (Figure 1 – Supplement 1D).

As expected, control RNAi did not result in any ectopic P-EN neurons, although there were a few neurons labeled outside the central brain and a small pattern of fan-shaped body neurons (Figure 6A-A’’’). In contrast, Eyeless^RNAi^ specifically in old INP progeny resulted in the formation of sparse populations of “late-born” ectopic P-EN neurons with projections into the PB, EB, and Noduli (Figure 6B-B’’’). These are the same neuropils targeted by wild type early-born P-EN neurons. We conclude that ectopic late-born P-EN neurons have morphology indistinguishable from the normal early-born P-EN neurons (Movies 1-2).

**Figure 6.**
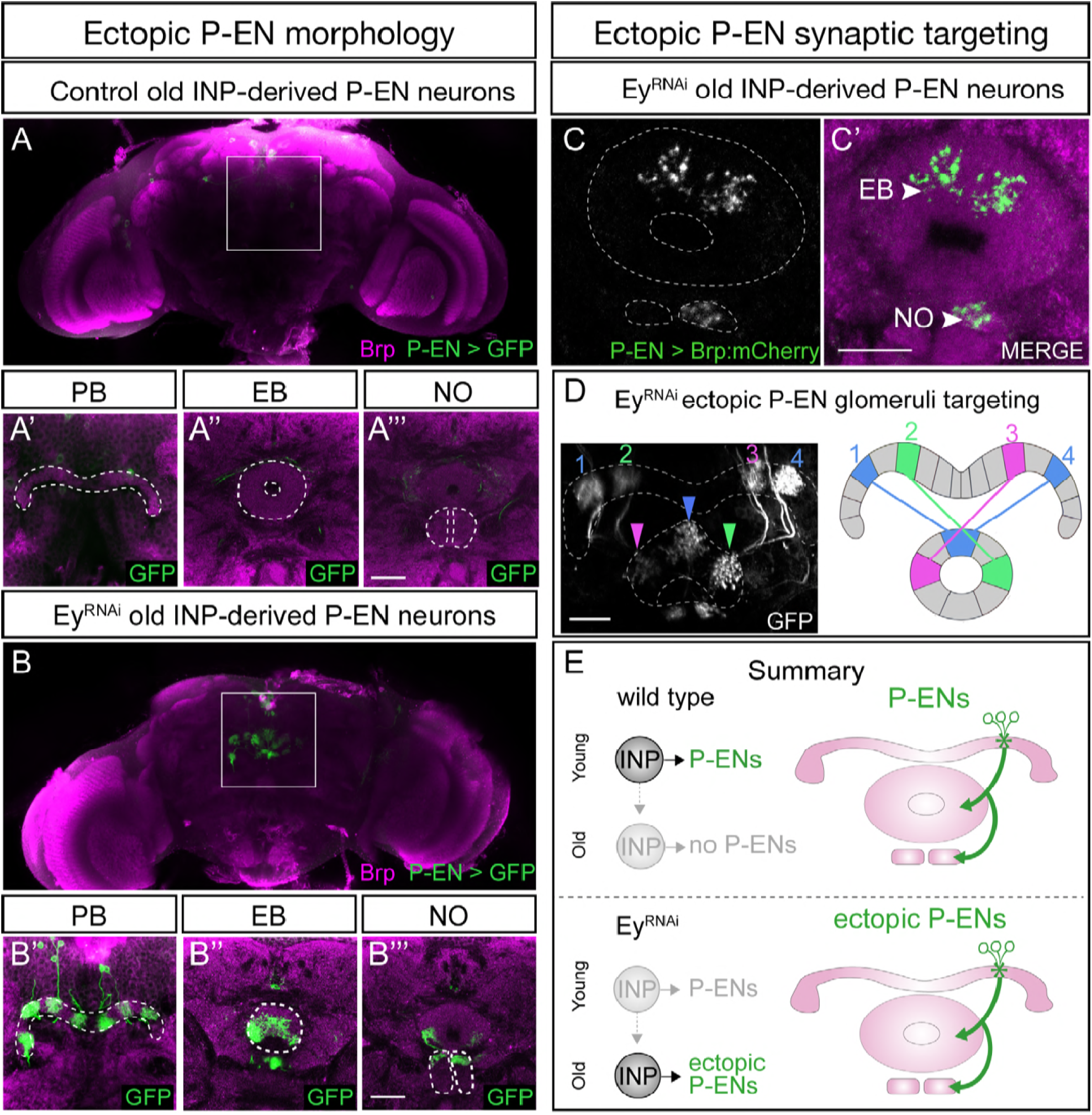
Eyeless^RNAi^ produces temporally transformed ectopic P-EN neurons that have normal morphology and connectivity. (A-A’’’) In wild-type adults, late INP clones do not label P-EN neurons in the adult brain (n=5). See methods for details. PB, EB, and NO neuropils marked with dashed lines. Scale bars, 20μm (A-B). (B-B’’’) In Eyeless^RNAi^ adults, late INP clones produce ectopic late-born P-EN neurons, which project to the PB, EB, and Noduli (n=5), similar to endogenous P-EN neurons (Figure 1A,B). See methods for details. PB, EB, and NO neuropils marked with dashed lines. (C-C’) In Eyeless^RNAi^ adults, late INP clones produce ectopic late-born P-EN neurons, which localize the pre-synaptic marker Brp::mCherry to the EB and Noduli (n=5), but not the PB (not shown), similar to the endogenous P-EN neurons. Scale bars, 20μm (C-D). Eyeless^RNAi^ adult, showing stochastic labeling of four ectopic P-EN neurons (1-4) with normal PB and EB glomeruli targeting (compare to Wolff et al., 2018 Figure 8D1). See methods for details. (E) Summary showing full temporal transformation of ectopic P-EN neurons following Eyeless^RNAi^.

To determine if the ectopic P-EN neurons have the same anatomical connectivity as the endogenous P-EN neurons, we expressed the pre-synaptic active zone marker Bruchpilot (Brp) specifically in the ectopic P-EN neurons. We found that ectopic P-EN neurons localized Brp to the EB and Noduli, but not to the PB. This is the precisely the same as wild type P-EN neurons (Figure 6C-C’, summarized in Figure 6E). Thus, the ectopic P-EN neurons match the normal early-born P- EN neurons in molecular identity (R12D09-LexA+), morphology (PB, EB, Noduli projections), and anatomical connectivity (Brp puncta in EB and Noduli). Finally, we determined that ectopic P-EN neurons assemble into proper columns between glomeruli in the PB and tiles in the EB, precisely matching the morphology of endogenous P-EN neurons (Figure 6D; compare to Wolff et al., 2015, Figure 8D1). We conclude that (1) reducing expression of the TTF Eyeless leads to a doubling of P- EN neurons in the CX, and (2) temporal identity determines P-EN morphology and anatomical connectivity, not time of neuronal birth or differentiation.

### Transient Eyeless reduction impairs adult flight navigation behavior

Our finding that the temporal transcription factor Eyeless contributes to the development of CX columnar neurons raises the question of how Eyeless influences CX function. Recent work has shown that silencing adult E-PG neurons impairs flies’ capacity to maintain an arbitrary heading to a bright spot resembling the sun (Giraldo et al. 2018; Green et al. 2018), a finding that we independently confirmed (Figure 7 – supplement 1A-D). Based on these results, we hypothesized that Eyeless function during development may be required for adult E-PG function in sun navigation. To reduce Eyeless expression, we drove Eyeless^RNAi^ in old INPs using *R16B06-Gal4*.

**Figure 7.**
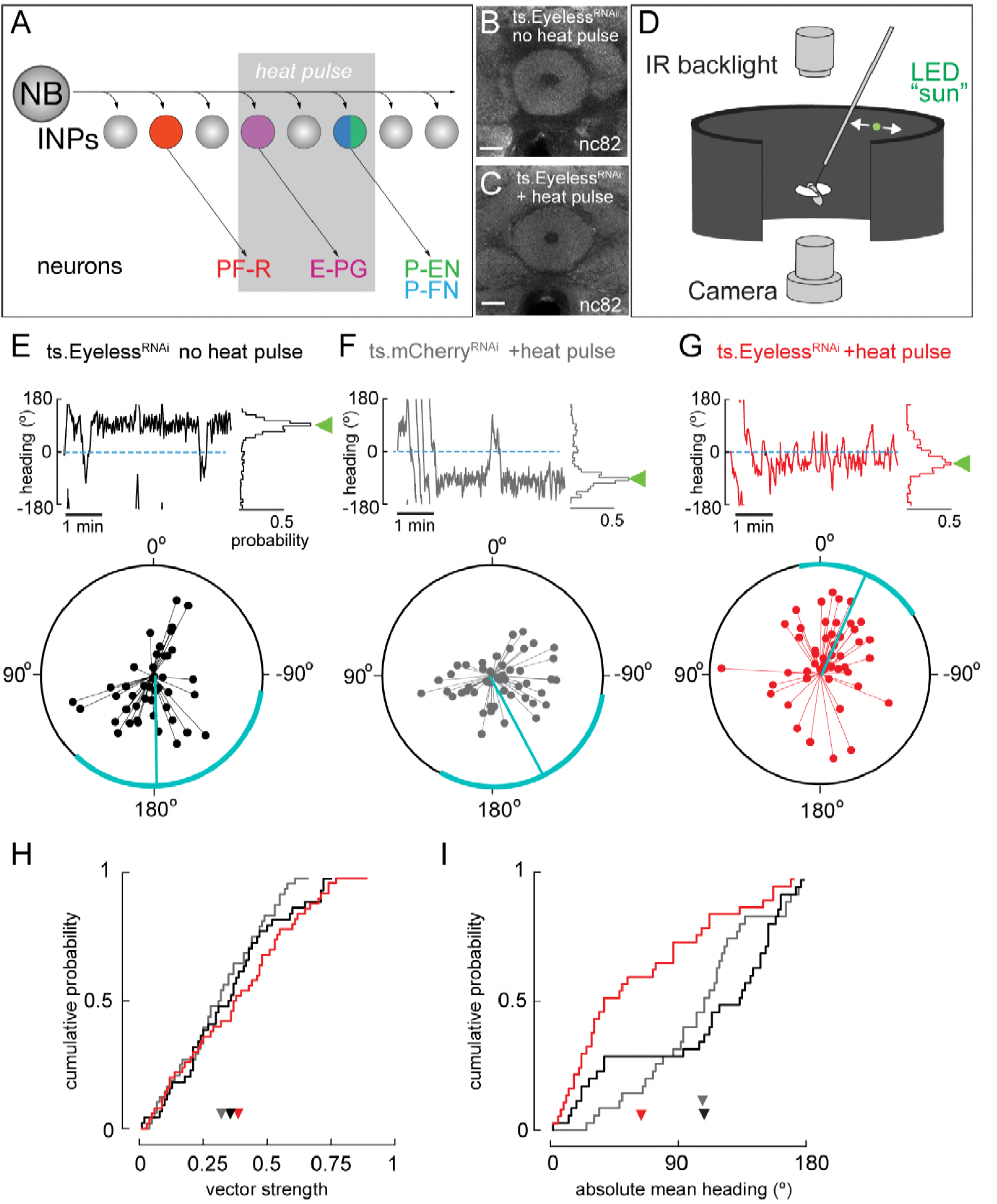
Transient loss of Eyeless during development impairs adult fly navigation. (A) Timing of Eyeless reduction in INP lineages. Transient inactivation of ts.Gal80 (29°C heat pulse; gray bar) results in transient Eyeless^RNAi^ during the time in which E-PG neurons are normally generated. (B,C) The manipulation in A does not alter CX neuropil morphology as seen by nc82 (neuropil) staining (EB, shown; other neuropils, data not shown). (D) Schematic of experimental apparatus for sun navigation experiments. The wing stroke amplitudes of a tethered, flying fly were monitored with an IR camera; the stroke difference determined the angular velocity of a 2.4° sun stimulus. Modified from (Giraldo et al. 2018). (E) Example flight (top panel) and summary data (bottom panel) from ts.Eyeless^RNAi^ control with no heat pulse. Top panel: left plot shows headings over 5 min flight; 0° is sun position in front of fly. Right histogram is distribution of headings in this example flight; sideways green triangle is the mean. Bottom panel: summary data, with each 5 min flight represented by radial lines. The angle of each line is the mean flight heading. The length of each line is vector strength of flight, varying from 0 (center of circle; no stimulus stabilization) to 1 (edge of circle; perfect stabilization). Each fly flew for two 5 min flights separated by a 5 min rest period. Cyan line shows mean heading, across flights with vector strength>0.2, as well as 95% confidence interval, calculated via resampling across flies. 44 flights, 22 flies total. (F) Example and summary data from ts.mcherry^RNAi^ control. Same plotting convention as (E). 48 flights, 24 flies. (G) Example and summary data from ts.Eyeless^RNAi^ flies with heat pulse. Same plotting convention as (E,F). 50 flights, 25 flies. (H) Cumulative probability distribution of vector strengths from both control groups (black and gray) and from experimental group (red). There was no significant difference between means (ts.Eyeless^RNAi^ heat pulse, 0.34, red; ts.Eyeless^RNAi^ no heat pulse, 0.31,black;ts.mcherry^RNAi^ heat pulse 0.30,gray; p>0.1, permutation test). (I) Cumulative probability distribution of mean absolute headings. The heading distribution for the experimental group, ts.Eyeless^RNAi^ heat pulse, was skewed significantly to frontal headings (mean 63.7°, 37 flights in 23 flies with vector strength>0.2) compared to control distributions (p<0.01, permutation test; ts.Eyeless^RNAi^ no heat pulse, mean 107.9°, 35 flights in 19 flies; ts.mcherry^RNAi^, mean 106.9°, 31 flights in 21 flies). Scale bars, 10μm.

Temporal control over Eyeless^RNAi^ was achieved with the temperature-sensitive Gal4 inhibitor Gal80. We raised animals at the Gal80 permissive temperature (18°C) to prevent Eyeless^RNAi^ expression and shifted to the non-permissive temperature (29°C) for 24h at the time E-PG neurons are born and differentiate (Figure 7A). Both control and Eyeless^RNAi^ animals exposed to this regime had no major morphological defects in the central complex (Figure 7B,C; additional data not shown). We then examined how the transient reduction of Eyeless in larval INPs affected the ability of adult flies to maintain an arbitrary flight heading to a fictive sun (Figure 7D). We compared the sun headings of Eyeless^RNAi^ flies that received the 29°C heat pulse with two control groups. One control group had an identical genotype but received no heat pulse (Figure 7E). A second control group received the heat pulse but Eyeless^RNAi^ was replaced with mcherry ^RNAi^ (Figure 7F). In both control groups, we found that flies maintained arbitrary headings, as expected, with a slight bias towards headings where the sun was behind the fly (Figure 7E,F,I). In contrast, flies with transient Eyeless^RNAi^ during E-PG development exhibited a marked frontal bias in their heading distribution, which was significantly more frontal than the control distributions (Figure 7G, I; p<0.01, permutation test). The control distributions were not significantly different from each other (p=0.49). Notably, although the heading distributions were distinct, the degree of stimulus stabilization – quantified by calculating the overall vector strength of each flight - was equivalent in the Eyeless^RNAi^ genotype and controls (Figure 7H). Moreover, the Eyeless^RNAi^ genotype and controls showed equivalent performance orienting to a dark vertical stripe (Figure 7 – supplement 1E-H), similar to the effect of silencing adult E-PG neurons (Giraldo et al. 2018; Green et al. 2018). This suggests that E-PG silencing and Eyeless^RNAi^ induce similar, relatively specific navigation deficits rather than a more general deficiency in visual-motor flight control. Taken together, our results indicate that a transient loss of Eyeless specifically in old INPs causes specific deficits in adult flight navigation to that of silencing E-PG neurons. Our findings therefore demonstrate the importance of Eyeless for CX function.

### The Eyeless target gene Toy is required for E-PG axonal connectivity to the Gall

The TTF Eyeless is required to specify E-PG neuronal identity, but Eyeless does not persist in adult E-PG neurons, raising the question: What Eyeless target genes regulate E-PG connectivity and function? We focused on Twin of eyeless (Toy) which encodes a transcription factor whose expression is induced by Eyeless in old INPs (Bayraktar and Doe 2013) and is maintained in their adult post-mitotic neuronal progeny. We used two previously characterized Gal4 drivers (Kim et al. 2017; Lovick et al. 2017) to express *UAS-toy*^*RNAi*^ specifically in post-mitotic E-PG neurons at different stages in development and confirmed that it removes all detectable Toy protein (data not shown).

We next determined if depleting Toy in post-mitotic larval E-PG neurons using *R19G02-Gal4 UAS-toy*^*RNAi*^ altered E-PG survival or morphology. Loss of Toy had no effect on E-PG neuronal number (n=5, p=0.92) or on connectivity to the EB and PB (data not shown). In contrast, we observed greatly diminished E-PG axonal connectivity to the Gall, where in some cases the E-PG projections appeared nearly absent (n=12, Figure 8A-C). We next removed Toy later, beginning ∼24h after pupal formation using *ss00096-Gal4 UAS-toy*^*RNAi*^, and observed no effect on E-PG neuronal number (n=5, p=.48) or projections to the EB, PB, or Gall (n=6, Figure 8D-F). Surprisingly, however, loss of Toy produced a significant reduction in the levels of the pre-synaptic active zone marker Bruchpilot (Brp) in the Gall (Figure 8G-I). We conclude that Toy is required during larval stages for E-PG connectivity to the Gall, and is required in pupal stages for establishing or maintaining Brp levels at the E-PG axonal terminals in the Gall.

**Figure 8.**
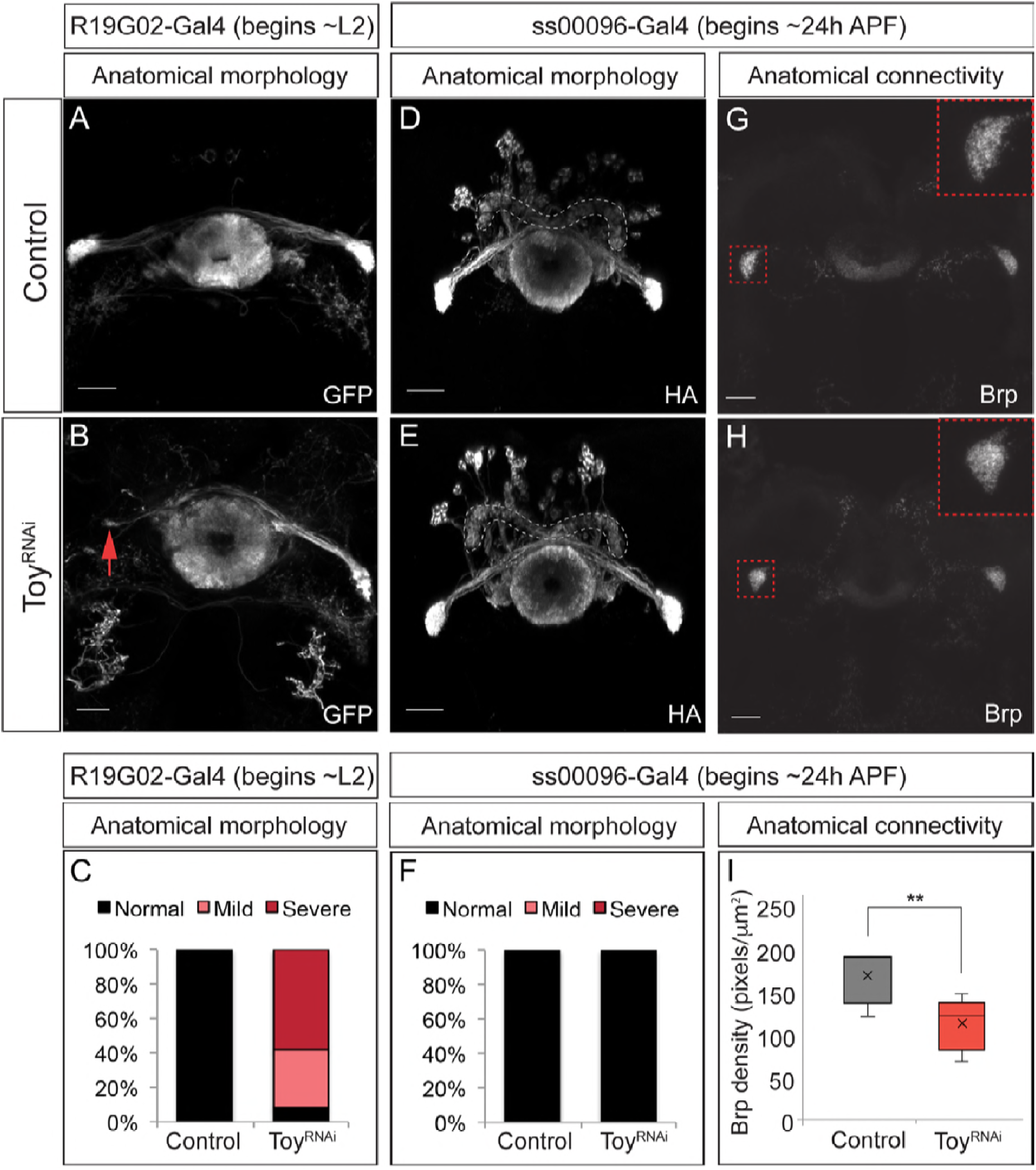
The Eyeless target gene Toy is required for E-PG axonal connectivity. (A-C) Loss of Toy in larvae reduces E-PG projections to the Gall in adults. (A) Wild type: *R19G02-Gal4* is first expressed at ∼L2 and labels adult E-PG neurons; note projections to the EB (center) and Gall (left and right); PB, not shown (n=6). (B) *R19G02-Gal4 UAS-Toy*^*RNAi*^ reduces E-PG projections to the Gall (red arrow), yet projections to the EB and PB (not shown) remain intact (n=12). (C) Quantification. (D-F) Loss of Toy in pupae has no effect on E-PG projections. (D) In wild type, *ss00096 split-Gal4* is expressed ∼24h after pupal formation and labels adult E-PG neurons; n=5. (E) *ss00096 split-Gal4 UAS- Toy*^*RNAi*^ adults have normal projections to the EB (center), Gall (left and right), and PB (outlined); n=5. (F) Quantification. (G-I) Loss of Toy in pupae reduces pre-synaptic levels of Brp in the Gall. (G,H) Genotypes as in D-E, showing that the pre-synaptic marker Brp is reduced in the E-PG axons targeting the Gall following ToyRNAi (n=6). (I) Quantification. Scale bars, 20μm.

To determine how the loss of Toy during pupal stages affects CX function, we tested whether reduction of Toy in the E-PGs affected sun navigation. We observed no significant change in flies’ heading distribution in relation to the sun stimulus, or in the degree to which they stabilized the sun stimulus (Figure 8 – Supplement 1). Therefore, the loss of Toy in pupal INPs and the associated reduction of Brp at E-PG axon terminals has no discernible effect on sun navigation.

## Discussion

### Developmental origin of CX columnar neurons

We have shown that distinct classes of CX columnar neurons have unique developmental origins within type II neuroblast lineages. We find that CX columnar neurons map to four bilateral type II neuroblast lineages (DM1-DM4), confirming previous work (Wang et al. 2014). Our birth-dating results indicate that CX columnar neurons originate from distinct INPs born ∼12h apart during larval life, except for P-EN and P-FN neurons whose similar birthdates suggest they may arise from the same INPs. Twin-spot MARCM analysis (Lee and Luo 1999) would be necessary to determine whether P-EN and P-FN neurons arise from the same or different INPs. Interestingly, the two CX columnar neurons born at the same time (P-EN and P-FN) have axon projections intrinsic to the CX and target the same neuropils (PB and Noduli). In contrast, the two CX columnar neuron types born at different times (E-PG and P-FR) have axon projections extrinsic to the CX and target different neuropils (Gall and ROB). This raises the possibility that neuroblast temporal identity determines whether columnar neuron axon projections are intrinsic or extrinsic to the CX. More generally, the results suggest that neurons with similar temporal identity have matching connectivity.

We have mapped the birthdates of only four CX columnar neuron subtypes out of the 60 distinct neuronal subtypes innervating the CX (Young and Armstrong 2010). Mapping these other neurons to their type II neuroblast and INP lineages is an important task for the future, which will help identify developmental correlates of neuronal morphology, connectivity, and function. Additionally, significant neuronal diversity may arise from GMCs dividing to make Notch^ON^/Notch^OFF^ sibling neurons, which often have distinct morphology (Truman et al. 2010; Lacin et al. 2014; Wang et al. 2014; Harris et al. 2015). The role of Notch signaling in generating hemilineages within type II neuroblast progeny remains unexplored.

### Specification of CX columnar neurons

By mapping the developmental origins of four classes of columnar neurons innervating the central complex, we find that each class derives from a relatively tight window during the neuroblast lineage, and from either young or old INPs (Figure 3I). The fact that all of the four subtypes are restricted to early or late in the INP lineage suggests that the early/late lineage distinction is developmentally important, consistent with our finding that early/late INPs express different TTFs (Dichaete/Eyeless, respectively). Furthermore, mapping the lineage of each neuronal class allowed us to identify a correlation with developmental origin and neuronal morphology (neurons with similar birth-dates have similar morphology). Many other developmental windows have yet to be characterized, for example the neurons derived from young INPs prior to PF-R/E-PG production are unknown, and would be expected to be expanded in the absence of Eyeless; similarly, the neurons derived from the old INPs following production of the P-EN/P-FN neurons are unknown, and would be expected to be missing in the absence of Eyeless. In the future, our intersectional genetic approaches can be used to map the developmental origin of any neuronal subtype for which there exists an adult LexA driver line. For example, we have recently mapped the CX dorsal fan-shaped body “sleep neurons” (Donlea et al. 2011; Ueno et al. 2012; Dubowy and Sehgal 2017; Donlea et al. 2018) to an old neuroblast developmental window (M. Syed, LS, and CQD, unpublished).

We have shown that Eyeless maintains a balance of early-born P-EN/P-FN neurons and late-born E-PG/PF-R neurons by triggering a switch from early-born to late-born neuronal identity. Loss of Eyeless generates fewer E-PG neurons and more P-EN neurons (Figures 4,5). We document the loss of late-born E-PGs here, but many other uncharacterized neurons are also likely to be lost, except during our heat pulse experiments where we tried to specifically target E-PG neurons (Figure 7). Similarly, we document the production of ectopic P-EN neurons in the absence of Eyeless, but many other early-born neuron populations are likely to be expanded. We considered performing clonal analysis to identify the neurons sharing an INP lineage with our four neural subtypes, but decides against it because INPs make morphologically different neurons at each division (Wang et al. 2014); we would not be able to map these neurons to early or late in the INP lineage, nor would we have molecular or genetic markers for these neurons. Determining the identity and birth-order of neurons within each INP lineage will be a difficult task for the future. Developing markers for the remainder of the 60+ different CX neuronal subtypes will be needed understand the breadth of Eyeless function in generating CX neuronal subtypes. Additional neuronal subtype markers will also be important to test the role of type II neuroblast candidate TTFs (Ren et al. 2017; Syed et al. 2017). We predict that at least some of these candidate TTFs will be required to specify the identity of the four columnar neuron classes described here.

### Determinants of connectivity in CX columnar neurons

We have shown that the ectopic P-EN neurons formed due to reduced Eyeless levels have morphology and anatomical connectivity that matches the endogenous P-EN neurons (i.e. Brp+ neurites to the EB and NO, and Brp-negative neurites to the PB)(Figure 6). It is unknown, however, whether these ectopic P-ENs are functionally connected to the normal P-EN circuit partners. This could be resolved through functional imaging experiments testing whether ectopic P-ENs receive the innervation from E-PG or delta7 neurons like endogenous P-ENs (Franconville et al. 2018) or whether they form functional inputs to known E-PG downstream neurons (Lin et al. 2013; Green et al. 2017; Turner-Evans et al. 2017). Furthermore, we demonstrate that the Eyeless target gene Toy is required for E-PG axonal connectivity to the Gall. Future work could elucidate the target genes of Toy through RNA-seq that are required for assembling this connectivity, such as downstream cell surface molecules, thus linking INP temporal identity to a direct mechanism for neuronal connectivity in a highly conserved adult brain region.

### The effects of Eyeless manipulation on navigation behavior

We found that reducing Eyeless expression during early development (24-48 h after larval hatching) causes a profound shift in how flies orient their flight relative to a fictive sun stimulus. Whereas control populations adopt a broad set of headings, with a slight bias for orientations where the sun is behind (Figure 7E,F,I), Eyeless^RNAi^ flies choose flight directions where the sun is in front (Figure 7G, I). A similar shift to a more frontal heading distribution occurs when E-PG neurons are silenced, either following expression of the Kir2.1 inward rectifying channel (Figure 7 – Supplement 1; Giraldo et al., 2017) or with a synaptic transmission blocker in walking flies (Green et al., 2018). The consistent shift to a frontal heading after both E-PG silencing and Eyeless^RNAi^ suggests that Eyeless^RNAi^ affects navigation behavior via perturbation of E-PG neurons, although we cannot rule out an effect on unknown late-born neurons. Eyeless^RNAi^ causes no gross deformities in the CX, suggesting that only a subset of E-PGs were eliminated by Eyeless^RNAi^, as opposed to the severe EB defects following loss of all E-PG neurons (Xie et al. 2017); in contrast, genetic silencing likely affects all E-PG neurons (Giraldo et al. 2018). The fact that similar behavioral effects are induced by our more subtle Eyeless manipulation and E-PG silencing suggests that sun navigation is highly dependent on E-PG neuron activity. One difference between the behavioral effects of Eyeless^RNAi^ and E-PG silencing is the degree to which flies stabilize the sun stimulus. Whereas silencing E-PG neurons significantly reduces the overall vector strength, a measure of the heading consistency within a flight (Figure 7 – Supplement 1; Giraldo et al., 2017; Green et al., 2018), there is no such reduction in vector strength in Eyeless^RNAi^ flies (Figure 7H). This difference could be due to the more limited scope of the Eyeless manipulation or it could reflect some capacity of the adult CX to compensate for the larval developmental defect. Taken together, our findings demonstrate that a specific navigation behavior – arbitrary orientation to a sun stimulus – depends on the precise expression and function of the Eyeless TTF during larval development. These results raise the question of how other types of navigation depend on the development and function of other CX neuronal subtypes.

## Materials and Methods

### Fly strains

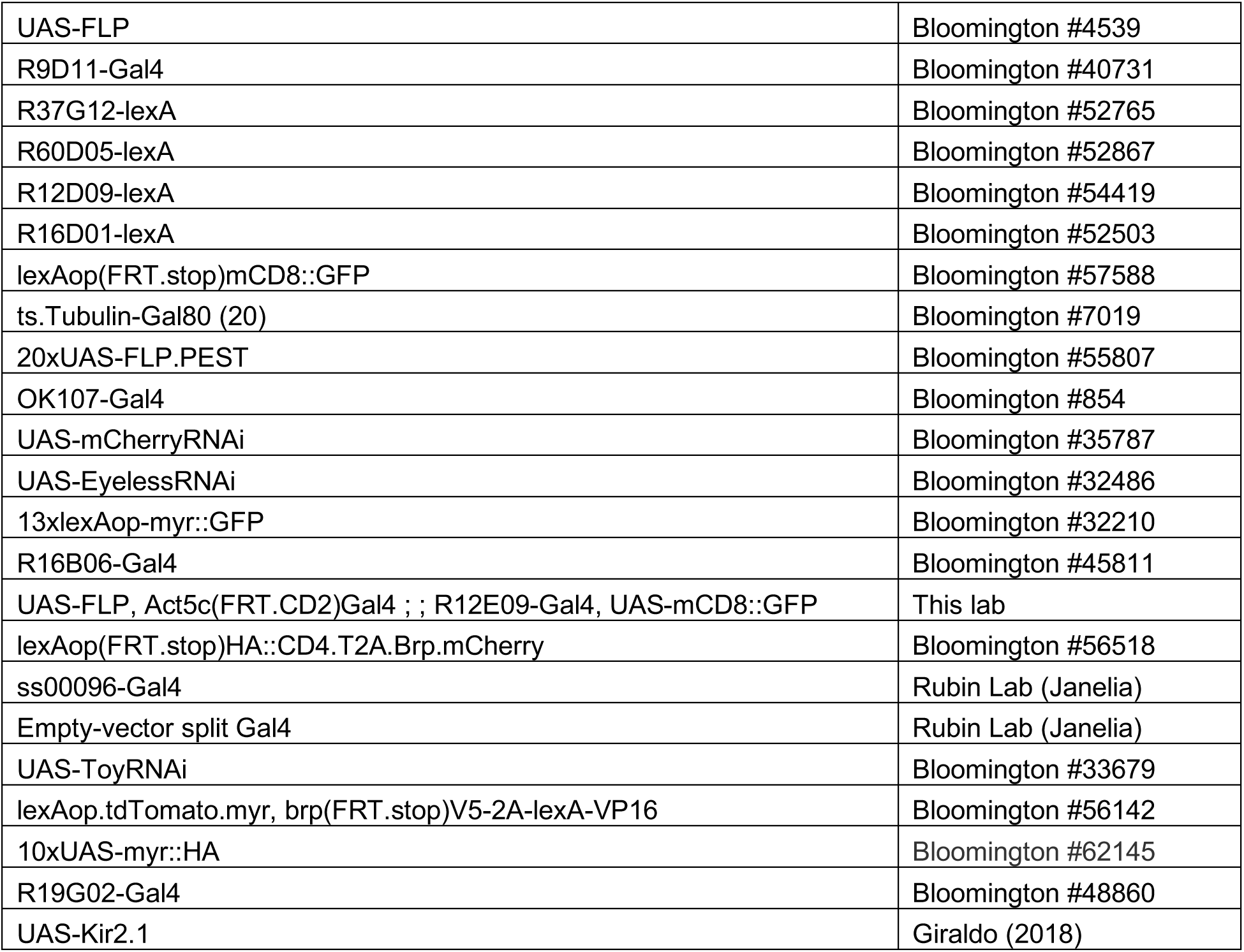

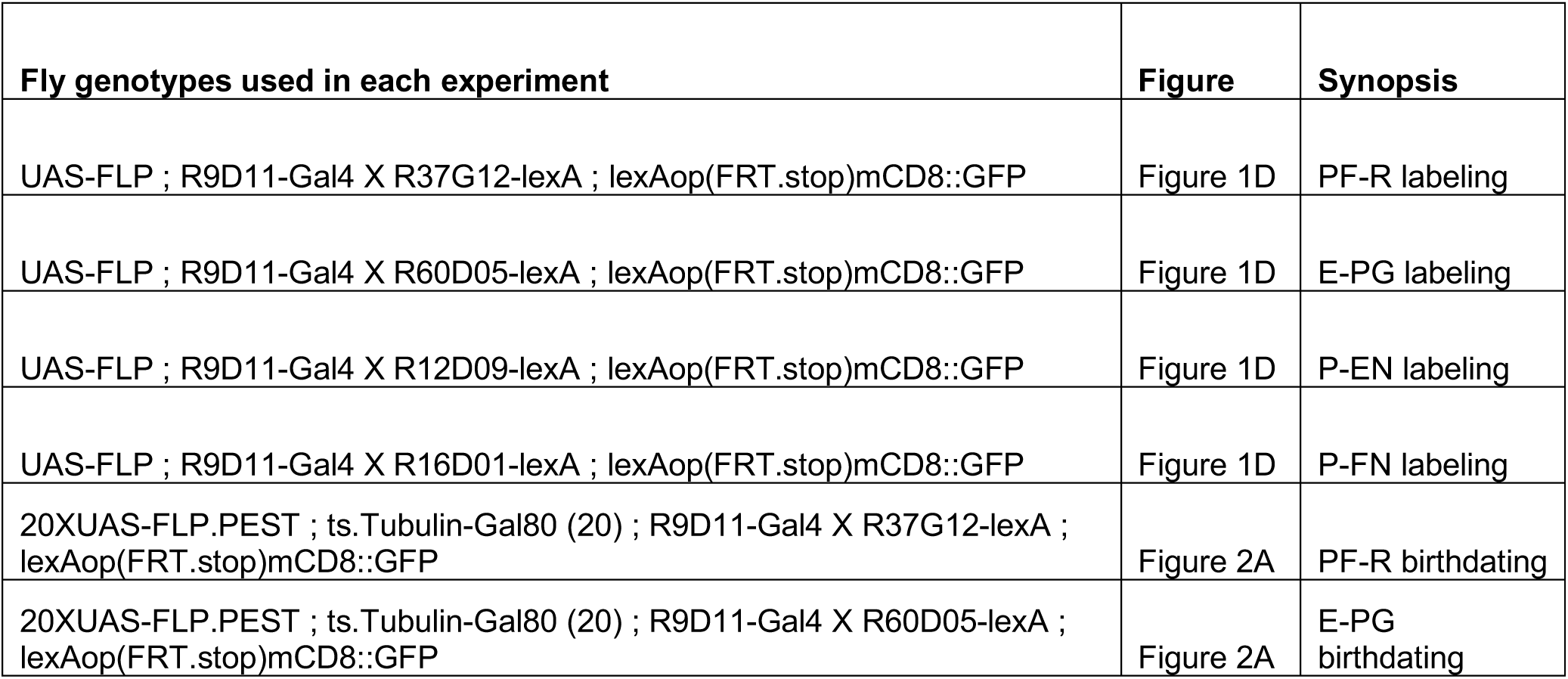
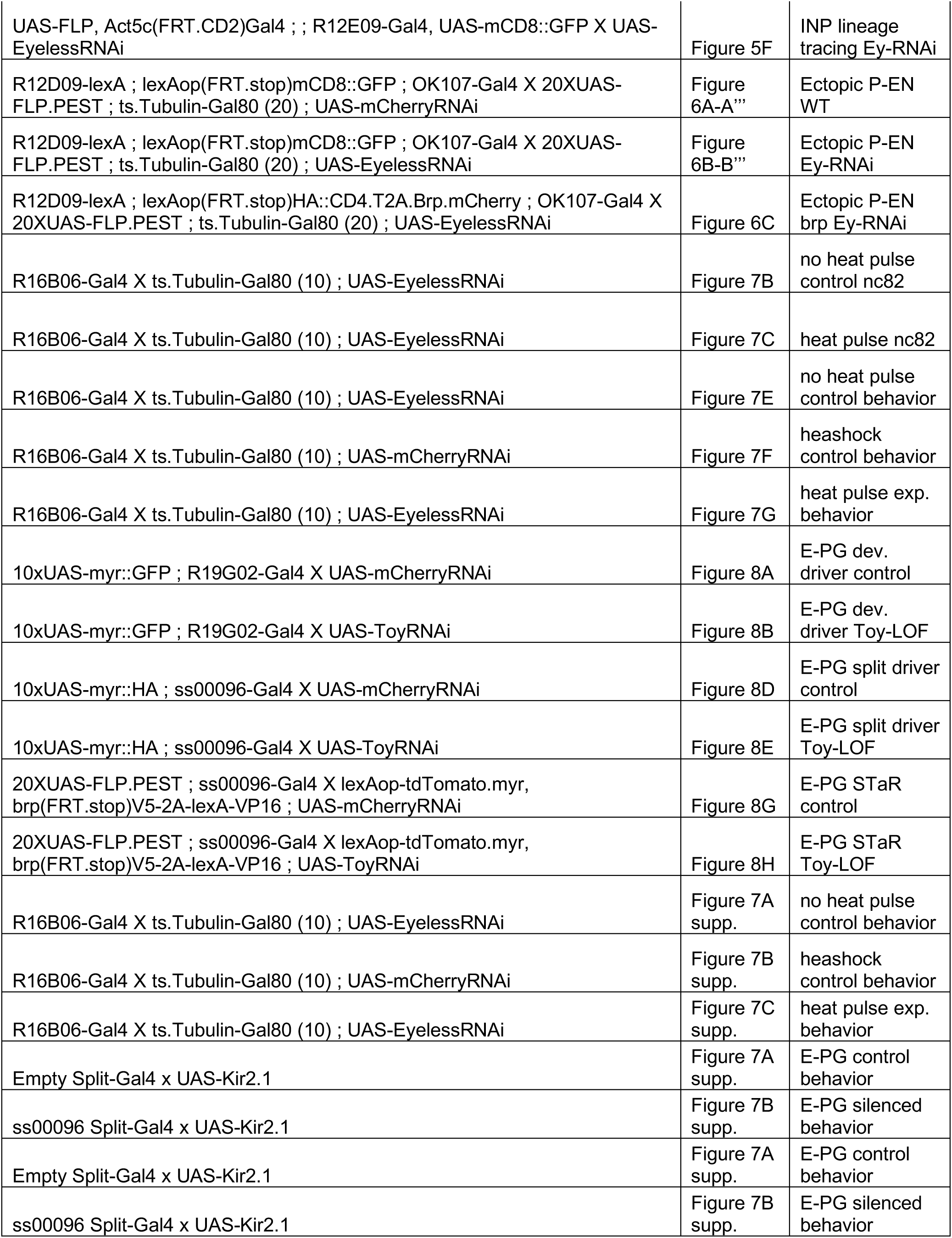
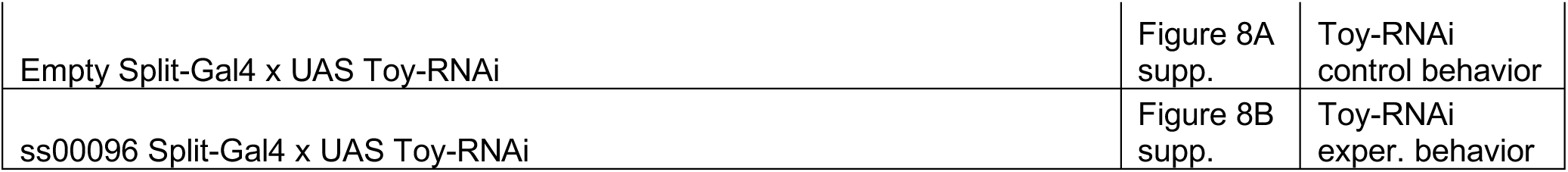

### Standardizing larval development at different temperatures

All larvae were grown at 25°C unless noted, and all hours after larval hatching are standardized to grow wild type at 25°C based on published conversions: 18°C is 2.25x slower than 25°C, and 29°C is 1.03x faster than 25°C (Powsner 1935).

### Immunohistochemistry

Primary antibodies were: chicken anti-GFP (1:1000, Abcam – Eugene, Oregon), rabbit anti-Toy (1:1000, Desplan Lab – NYU), guinea-pig anti-Runt (1:1000, Desplan Lab – NYU), and mouse anti- nc82 (1:50, DSHB – Iowa City, Iowa). Secondary antibodies were from Thermofisher or Jackson ImmunoResearch and were used at 1:400. Adult brain dissections were conducted at room temperature with 2-5 day old adult females. Adult brains were dissected in 2% formaldehyde solution in Phosphate-Buffered Saline with .5% Triton-X (PBST) and incubated for 55 minutes before applying an overnight block solution (5% Goat/Donkey serum, Vector Laboratories) at 4°C. Brains were then washed in PBST for one hour before applying an overnight primary mix at 4°C. Then, brains were washed for one hour at room temperature in PBST, before applying an overnight secondary mix at 4°C. Finally, brains were mounted in 90% glycerol, and imaged immediately.

### Imaging, data acquisition, and image analysis

Fluorescent images were acquired on a Zeiss LSM 700. Adult brain cell counting was performed using the FIJI cell counter plug in, and statistical analysis (Student’s T test) was done in Excel. Figures were assembled in Illustrator (Adobe). Relative Brp-density was quantified in FIJI; Slices were summed together in a z-stack, a rectangular ROI selected around the Gall -area, and a histogram plot of pixel intensity was generated. Background for image was calculated in neighboring ROIs and subtracted from each individual histogram plot-value. Intensity values were then summed together to calculate total intensity, and this was divided by Gall total area, calculated manually in FIJI using polygon selection tool.

### Fly tethering for flight behavior

We used 3-4 day old females for behavioral experiments. We tethered flies under cold anesthesia, gluing a tungsten wire to the anterior notum with UV-cured glue (Bondic). The head was immobilized relative to the body with a small amount of glue between the head and thorax. Flies recovered for at least 20 min prior to behavioral testing.

### Flight Arena and behavioral protocol

We coupled the angular velocity of a visual stimulus that was presented via LED panels to the continuously measured difference in wing stroke amplitude. Stroke amplitude was tracked at 60 Hz via Kinefly, a previously described video tracking system (Suver et al, 2016). A digital camera equipped with macro lens (Computar MLM3x-MP) and IR filter (Hoya) captured wing images from a 45° mirror positioned beneath the fly. Backlit illumination of wings was provided by a collimated infrared LED above fly (Thorlabs #M850L3). We displayed visual stimuli using a circular arena of 2 rows of 12 LED panels (24 panels total). Each panel had 64 pixels (Betlux #BL-M12A881PG-11, λ=525 nm) and was controlled using hardware and firmware (IORodeo.com) as previously described (Giraldo ref.).The gain between stimulus angular velocity and wing stroke amplitude difference was 4.75°/s per degree of wing stroke difference. The sun stimulus was a single LED pixel (∼2.4° on fly retina; Giraldo), ∼ 30deg above fly. The stripe was 4 pixels wide and 16 pixels high (15° by 60°). Flight experiments were controlled by custom scripts in the ROS environment. Incoming video was collected at 60 Hz and stimulus position data (i.e. the flight heading) at 200 Hz. In each experiment, flies navigated in closed loop to the sun stimulus in two distinct 5 min trials, which were separated by a 5 min rest period, during which we gave flies a small piece of paper to manipulate with their legs. Following the second sun flight, flies flew for 5 min in closed loop to the stripe stimulus. We discarded flights in which a fly stopped flying more than once during a sun or stripe presentation; furthermore, we discarded flights from flies that did not complete the two 5 min sun flights.

### Behavioral Data Analysis

All data analysis was conducted using custom scripts in Python. The circular mean heading of a flight was computed as the angle of resultant vector obtained via vector summation, treating each angular heading measurement as a unit vector. To determine the vector strength, we normalized the length of the resultant vector by the number of individual headings.

### Statistics

Data represent mean +/- standard deviation. Two-tailed Student’s t-tests were used to assess statistical significance of anatomical data, with *P<0.05; **P<0.01; ***P<0.001. To determine the significance of differences in the mean of the vector strength and heading distribution between groups, we used Fisher’s exact test with 10,000 permutations (Fisher, 1937). To avoid pseudoreplication, we permuted across flies rather than flights. We computed a 95% confidence interval of the circular mean of each heading distribution by bootstrapping from the observed data. For each experimental condition, we resampled with replacement from the observed flight data (resampling across flies not flights) to create 10,000 distributions of matched size to the observed data set. Confidence intervals were computed from the circular means of these 10,000 distributions. For analysis of the heading distributions and confidence intervals, we considered flights with a vector strength above a minimum threshold of 0.2.

## Acknowledgements

We thank the Tanya Wolff and Gerry Rubin for fly stocks. We thank Michael Dickinson for support in conducting the behavioral experiments. We thank Mubarak Syed, Emily Sales, Tanya Wolff, Claude Desplan, John Tuthill, and Michael Dickinson for comments on the manuscript. We thank Cooper Doe for assistance with brain dissections. Stocks obtained from the Bloomington Drosophila Stock Center (NIH P40OD018537) were used in this study. Funding was provided by HHMI (CQD, LS, TLW), NIH R37HD27056 (LS), and NIH T32HD007348 (LS).

## Author Contributions

LS and CQD conceived of the project; LS collected data for Figures 1-6 and 8, while TLW collected data for Figures 7, 7S, and 8S; LS, TLW, and CQD wrote the paper and prepared the figures. All authors commented and approved of the manuscript.

---

**Figure 1 – supplement 1.**
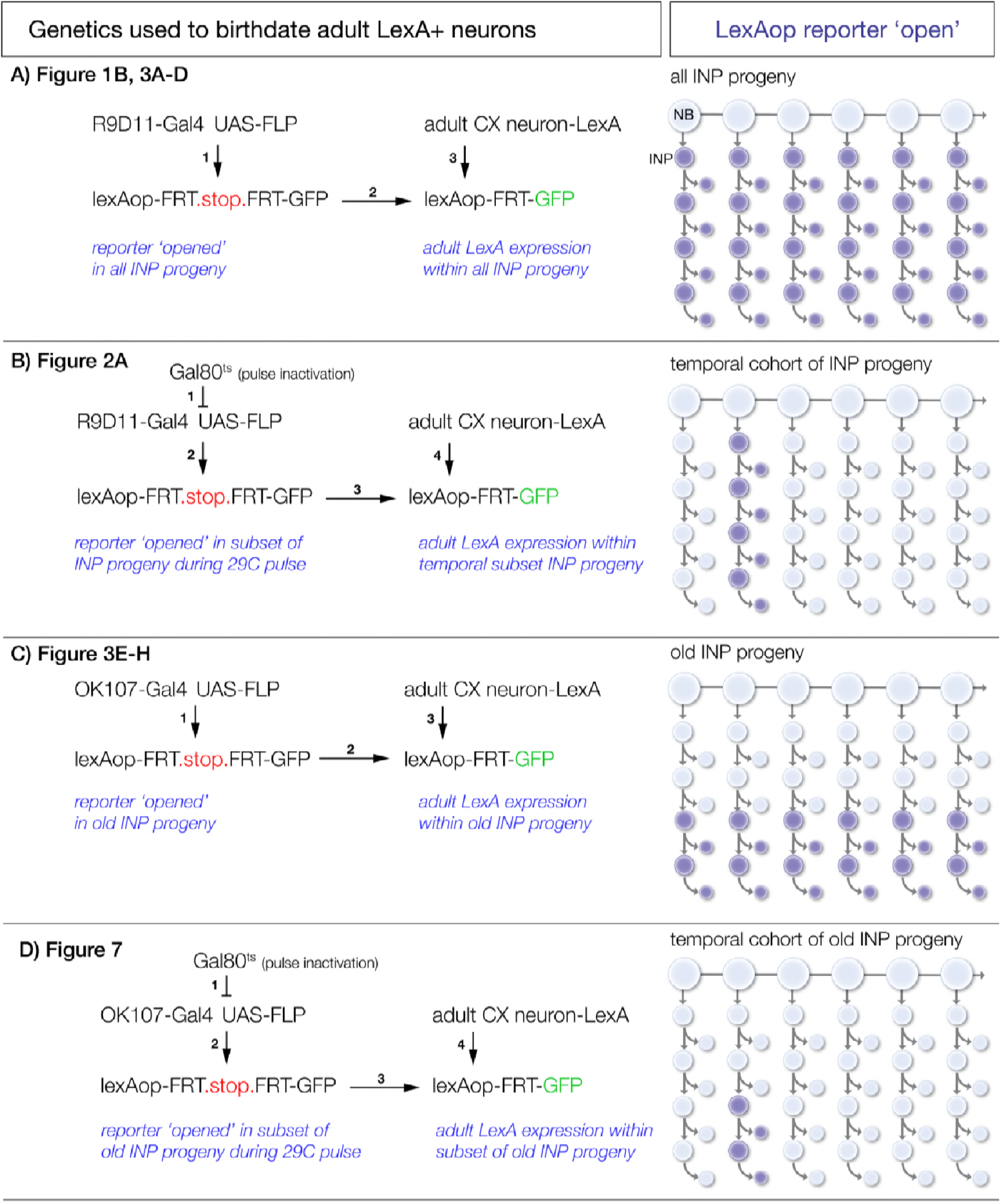
Intersectional genetic birthdating schemes. (A) Identifying columnar neuron subtypes derived from all INPs. *R9D11-Gal4* drives expression of FLP which excises an *FRT-stop* in all INPs and their progeny. This allows columnar neuron specific LexA lines to drive GFP expression if the neurons derive from INPs. (B) Identifying the time during the neuroblast lineage that produces each columnar neuron subtype. A pulse of 29°C disables ts.Gal80 to allow *R9D11-Gal4* to excise *FRT.stop* in the INPs present during the heat pulse. This allows columnar neuron specific LexA lines to drive GFP expression if the neurons derive from INPs born from the type II neuroblast at the time of heat pulse. (C) Identifying columnar neuron subtypes derived from young or old INPs. *OK107-Gal4* drives expression of FLP which excises an *FRT-stop* only in old INPs and their progeny. This allows columnar neuron specific LexA lines to drive GFP expression if the neurons derive from old INPs. Lack of expression shows the neurons are derived from young INPs. (D) Identifying the time during the neuroblast lineage that produces each old INP-derived columnar neuron subtype. A pulse of 29°C disables ts.Gal80 to allow *OK107-Gal4* to excise the *FRT.stop* in the INPs present during the heat pulse. This allows columnar neuron specific LexA lines to drive GFP expression if the neurons derive from INPs present at the time of heat pulse.

**Figure 7 – supplement 1.**
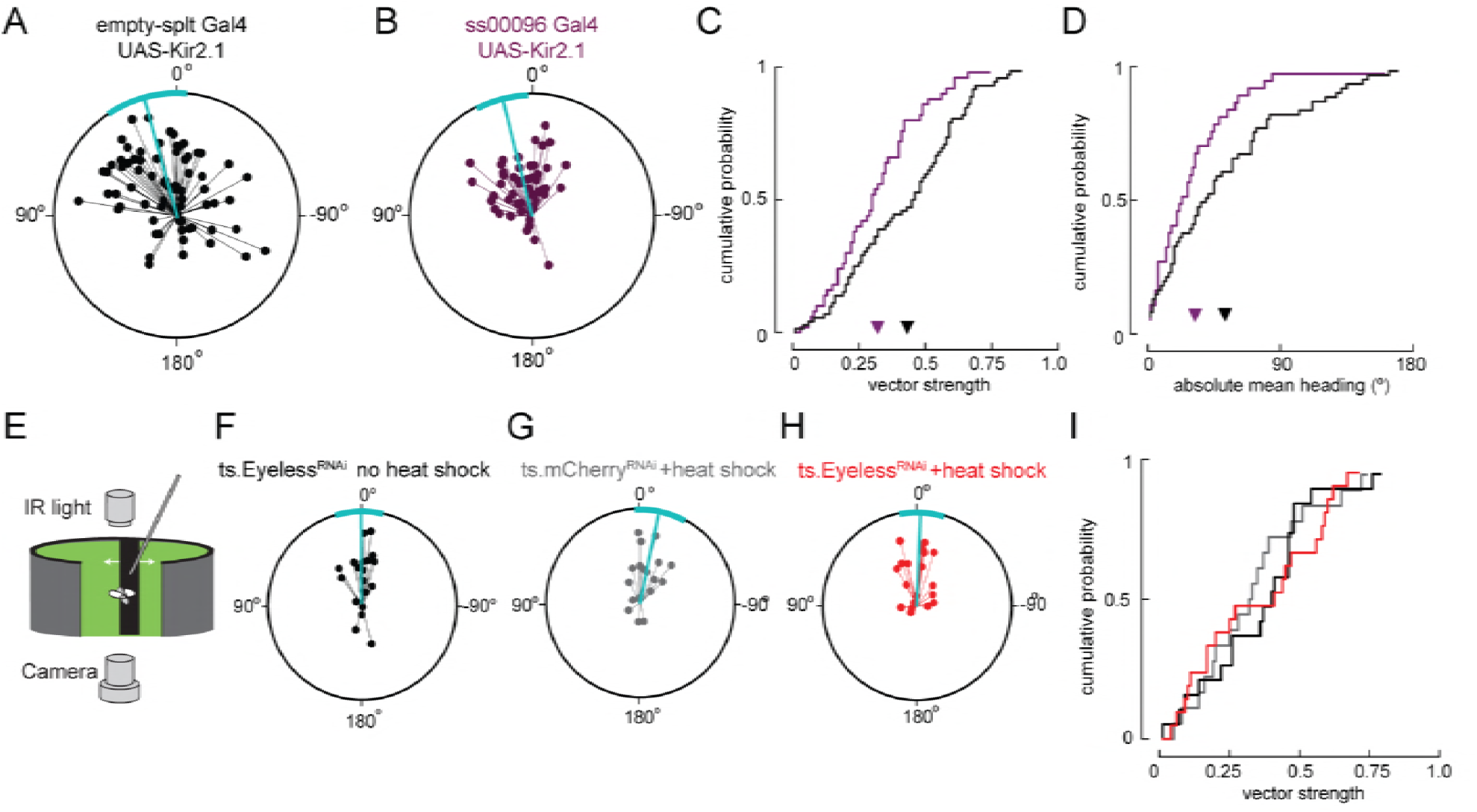
Sun navigation is impaired when E-PG neurons are silenced; stripe fixation behavior is not altered by the loss of Eyeless. (A-D) Silencing E-PG neurons impairs the flies’ capacity to maintain an arbitrary heading to a bright spot resembling the sun. (A) Flight headings to sun stimulus in *empty-Split / UAS Kir 2.1* control flies. Plotting convention same as Figure 7E-G. N=72 flights, 36 flies. (B) Flight headings to sun stimulus in *ss00096 Split-Gal4 (E-PG neurons) / UAS-Kir2.1* flies. N=50 flights, 25 flies. (C) Cumulative probability distribution of vector strength values for flights to sun stimulus. *empty Split-Gal4 UAS-Kir2.1* (black line), mean=0.43, N=72 flights, 36 flies; *ss00096 Split-Gal4 / UAS-Kir2.1* (purple line), mean=0.32, N=50 flights, 25 flies. P=.001, permutation test for significant difference in mean. (D) Cumulative probability distribution of mean absolute heading for flights to sun stimulus. *empty Split-Gal4 UAS-Kir2.1* (black line), mean=52.7°, N=61 flights, 36 flies; ss00096 *Split-Gal4 / UAS-Kir2.1* (purple line), mean=32.2°, N=37 flights, 23 flies. P=.011, permutation test for significant difference in mean. (E-H) Transient removal of Eyeless in larvae at the time of E-PG development (as in Figure 7) has no effect on stripe fixation behavior in adult flies. Plotting conventions as in Figure 7E-G. (E) Schematic of behavioral apparatus; flies controlled rotation of a dark vertical stripe with their wing strokes. (F) Control ts.Eyeless^RNAi^ no heat pulse. N=19 flies. Flight headings are tightly clustered around 0°, where stripe is in front of fly. (G) Control ts.mcherry^RNAi^ flies with heat pulse. N=18 flies. (H) Experimental ts.Eyeless^RNAi^ flies with heat pulse. Heading distribution remains centered around 0°. N=21 flies. (I) Cumulative probability distribution of vector strength values for stripe stimuli with control groups (black and gray) and experimental group (red). We observed no significant difference between means for all genotypes.

**Figure 8 – supplement 1.**
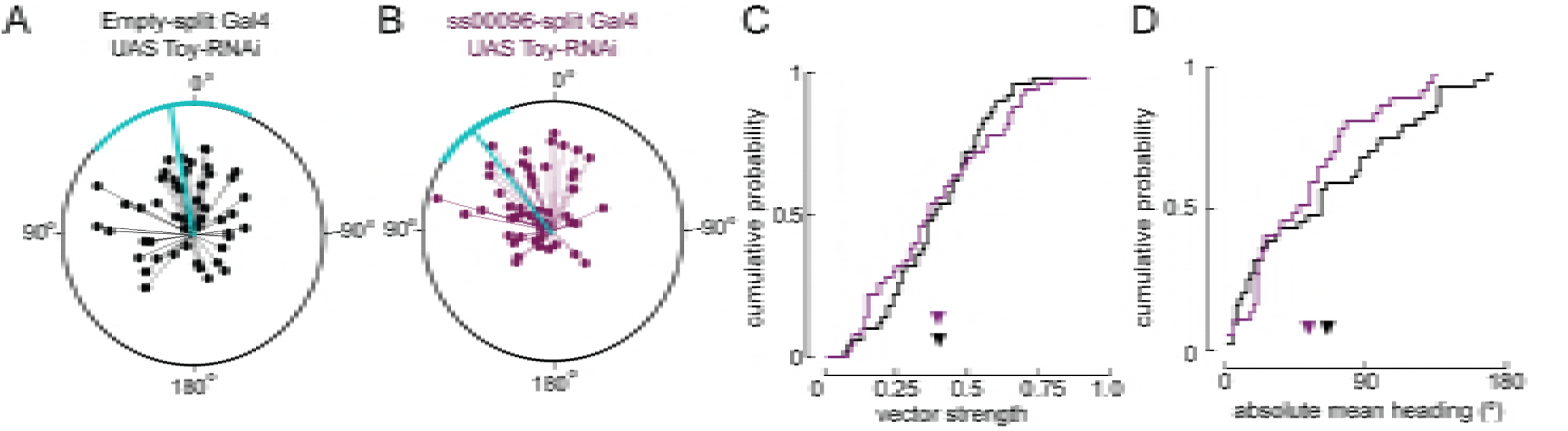
Reducing levels of Toy in E-PG neurons during pupal stages does not alter navigation to a fictive sun. (A) Flight headings to sun stimulus in *empty-split Gal4 / UAS Toy-RNAi* control flies. As in Supplementary Figure 7, each 5 min flight is represented with a distinct radial line. The angle of the line is the mean flight heading and the length of line is vector strength. The cyan line shows mean heading across flights with vector strength>0.2 as well as a 95% confidence interval, calculated via resampling across flies. N=72 flights, 36 flies. (B) Flight headings to sun stimulus in *ss00096-split Gal4 (E-PG neurons) / UAS Toy-RNAi* flies. Plotting convention as in (A). N=42 flights, 21 flies. (C) Comparison of cumulative probability distribution of vector strength values. Control *empty-split Gal4 UAS Toy-RNAi* (black line), mean=0.40; *ss00096-split Gal4 / UAS Toy-RNAi* (purple line), mean=0.40. (D) Cumulative probability distribution of mean absolute heading for flights to sun stimulus. *empty-split Gal4 UAS / Toy-RNAi* (black line), mean=65.4°, N=44 flights, 25 flies; pupal *ss00096-split Gal4 / UAS Toy-RNAi* (purple line), mean=53.2°, N=37 flights, 21 flies. Permutation test for significant difference of means, P=0.17.

**Supplemental Movie 1. wild-type adult P-EN morphology**

wild-type young INP born P-EN neurons innervate distinct neuropil regions of the central complex. These include the PB, EB, and NO outlined in red, white, and yellow respectively.

**Supplemental Movie 2. Ey-RNAi ectopic adult P-EN morphology**

Ey-RNAi ectopic old INP born P-EN neurons innervate the same distinct neuropil regions of the central complex. These include the PB, EB, and NO outlined in red, white, and yellow respectively. Note, only even-numbered glomeruli are generated, indicating they are born after odd numbered glomeruli in the INP lineage, and extend when Eyeless is eliminated in old INPs.

